# A refutation to ‘A new A-P compartment boundary and organizer in holometabolous insect wings.’

**DOI:** 10.1101/486951

**Authors:** Peter A. Lawrence, José Casal, José F. de Celis, Ginés Morata

## Abstract

We respond to a recent report by Abbasi and Marcus who present two main findings: first they argue that there is an organiser and a compartment boundary within the posterior compartment of the butterfly wing. Second, they present evidence for a previously undiscovered lineage boundary near wing vein 5 in *Drosophila*, a boundary that delineates a “far posterior” compartment. Clones of cells were marked with the *yellow* mutation and they reported that these clones always fail to cross a line close to vein 5 on the *Drosophila* wing. In our hands *yellow* proved an unusable marker for clones in the wing blade and therefore we reexamined the matter. We marked clones of cells with *multiple wing hairs* or *forked* and found a substantial proportion of these clones cross the proposed lineage boundary near vein 5, in conflict with their findings and conclusion. As internal controls we showed that these same clones respect the other two well established compartment boundaries: the anteroposterior compartment boundary is always respected. The dorsoventral boundary is mostly respected, and is crossed only by clones that are induced early in development, consistent with many reports. We question the validity of Abbasi and Marcus’ conclusions regarding the butterfly wing but present no new data.

Arising from: R. Abbasi and J. M. Marcus *Sci. Rep*. 7, 16337 (2017); https://doi.org/10.1038/s41598-017-16553-5

## Introduction

In many animals, single embryonic cells can be marked by genetic or other methods and their clonal progeny followed and mapped through later development. This approach has been extensively used in *Drosophila* and some other animals to provide information about patterns of growth and the serial acquisition of cell identity. One important discovery was that of developmental compartments, groups of cells that become determined early whose descendents subsequently fill precisely defined territories ^1–3^. These territories remain intact because the cells in each compartment appear to acquire differences in cell affinity that stop them mixing with cells of neighbouring compartments as they grow ^4^. Compartments are also units of gene function, their constituent cells as defined by lineage can also be the domains of action of a special class of genes, “selector genes”^5^. Examples are the *engrailed* gene that specifies and drives the affinity of all the cells of the posterior compartments in each insect segment ^6,4^ and *apterous*, that specifies the dorsal compartments of the wing and haltere ^7–10^.

The fly wing is made of four compartments; these are delineated by the anteroposterior (AP) compartment boundary that divides the wing into two halves and the dorsoventral (D-V) boundary that runs along the perimeter between the two epithelial surfaces. Also, these two boundaries align with sources that generate gradients of the two organiser molecules, Dpp and Wg ^11–16^. Our current understanding of the patterning processes in the wing is based on these two boundaries ^17–21^.

A recent report ^22^ has described aspects of wing development in both *Drosophila* and in butterflies. Abbasi and Marcus ^22^ report specific eyespot associations after examining wings of 22 different species of *Vanessa*, known to have highly diverse eyespot patterns. They describe correlations between eyespots that point to the existence of an additional organiser in the vicinity of vein M3 (note that use of this logic has been questioned by experiments performed by Beldade et al. ^23^). They also argue that, in *Drosophila*, the main organiser of the wing is associated with the A-P compartment boundary and conclude that the M3 organiser should also correspond with a boundary of cell lineage restriction. Strangely, the correlations they present as an argument for the M3 organiser (Table 1 in ref 22) do not detect the presence of the A-P organiser whose existence is independently established ^24^. Strange, because we would expect the A-P organiser to be detected at least as well as the F-P organiser; indeed, the A-P organiser should act as a positive control that should validate the method they used.

**Table 1.**
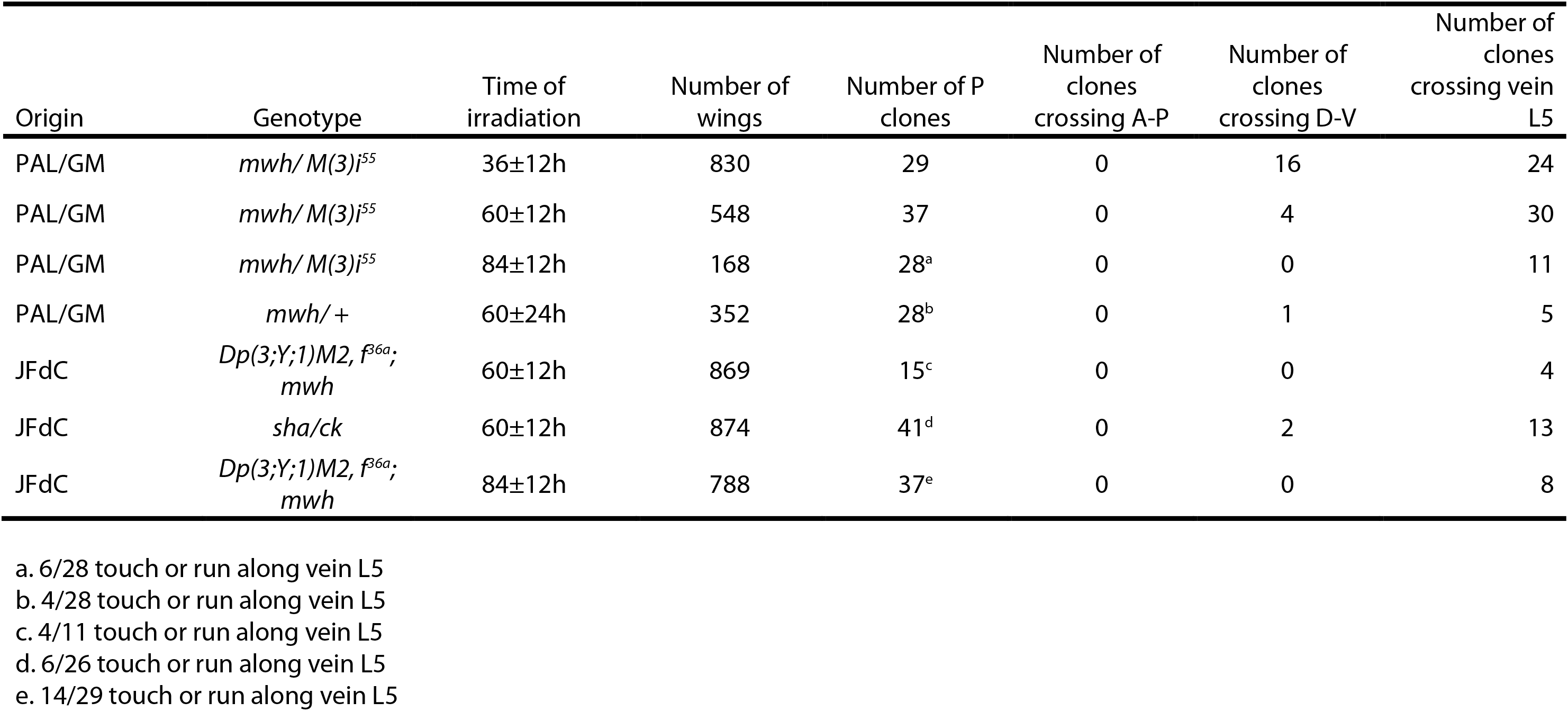

In parallel, their survey of naturally occurring mosaic butterfly wings (probably of different origins, including gynandromorphs) suggests the existence of a compartmental lineage boundary, that they call F-P for “far posterior”, also near to vein M3 although quantitative data supporting their conclusion is not presented. Studies on flies ^25–27^ have argued that quantitative data would be necessary to define a lineage boundary also in butterflies as many patches in gynandromorphs cross compartment boundaries — because their primordia arise from contiguous cells in the early embryo and because the male and female cells segregate long before the definition of the founding cells of compartments.

Abbasi and Marcus have marked clones in *Drosophila* wings to ask whether a similar lineage boundary may exist in flies. They claim it does —“Mitotic wing clones never cross an apparent Far-Posterior (F-P) compartment boundary just posterior of *Drosophila* vein L5, which is equivalent and homologous to vein M3 in butterfly wings”— and conclude this “finding of an additional, previously undocumented compartment boundary can be considered a breakthrough in the understanding of insect wing development and patterning”. Here we question the existence of this compartment boundary in *Drosophila*.

## Results and Discussion

In the 1970s, we and others studied cell lineage in the wing by making genetically marked clones rather as Abbasi and Marcus have done ^28–30,1,27^. In our experiments we used the mutations *multiple wing hairs* (*mwh*) and *forked, f*^36^, both of which can be scored cell by cell on both wing surfaces. Both these mutants are homozygous viable and have no significant effect on wing size, shape or pattern. Furthermore, cell lineage clones induced in *mwh* wings showed normal clone shapes and sizes, suggesting that this mutant does not affect the growth (clone size), mitotic orientation (clone shape), cell adhesion (clone integrity, clones do not split as they grow) during larval life ^31^. The clones were generated by X-rays and, in some experiments, we conferred the marked clones with a proliferation advantage so that they reveal their full potential ^32^. We find that clones induced in the embryo defined and respected the A-P compartment boundary and none crossed it. Clones induced early in larval development frequently crossed from dorsal to ventral surfaces on the wing blade but, following later irradiation, the clones always respected and defined that D-V boundary (Table 1). We found no other compartment boundaries. We (PAL and GM) have reexamined these preparations again to check for the existence of the F-P boundary. We find no evidence for such a boundary: the majority of early induced clones cross from one side to the other of the line delineating the proposed boundary. Even at the later time of clonal induction, when the dorsoventral boundary is respected by all clones, clones still very frequently cross over vein 5 and, as far as we can detect, respect no virtual line near to it (Table 1). We submit that the quantitative data in Table 1 also argue strongly against the existence of a F-P lineage boundary — even though the probability of two clones in the same wing could be higher than that suggested by the average numerical frequency of clones in the whole experimental population. In Table 1, we use independently established compartment boundaries as internal controls to check on the possibility that some of our “clones” could actually descend from two or more independent events. The Table documents that the A-P boundary is never crossed in our large data sets; which it should have been had some some of our clones had been due to double hits. The same argument relates to the D-V boundary which is crossed more when the clones are induced earlier than later, as would be expected. The data show that, when compared with either the A-P boundary or even the later established D-V boundary, the F-P boundary is more frequently crossed. These data argue against the proposed F-P compartmental boundary, even if it were established late in development.

The clones made by Abbasi and Marcus ^22^ did not have a growth advantage and this might explain the discrepancy between our results and theirs. We have therefore studied clones that had no growth advantage, being marked with *mwh*. Although Abbasi and Marcus ^22^ report that their clones “never” crossed the F-P boundary, a goodly number of ours did, crossing over vein L5 from both sides (Table 1, Figure 1); we confirm that some clones do tend to run along veins (including L5) for variable distances, as noted by Gonzalez-Gaitan et al ^33^.

**Figure 1.**
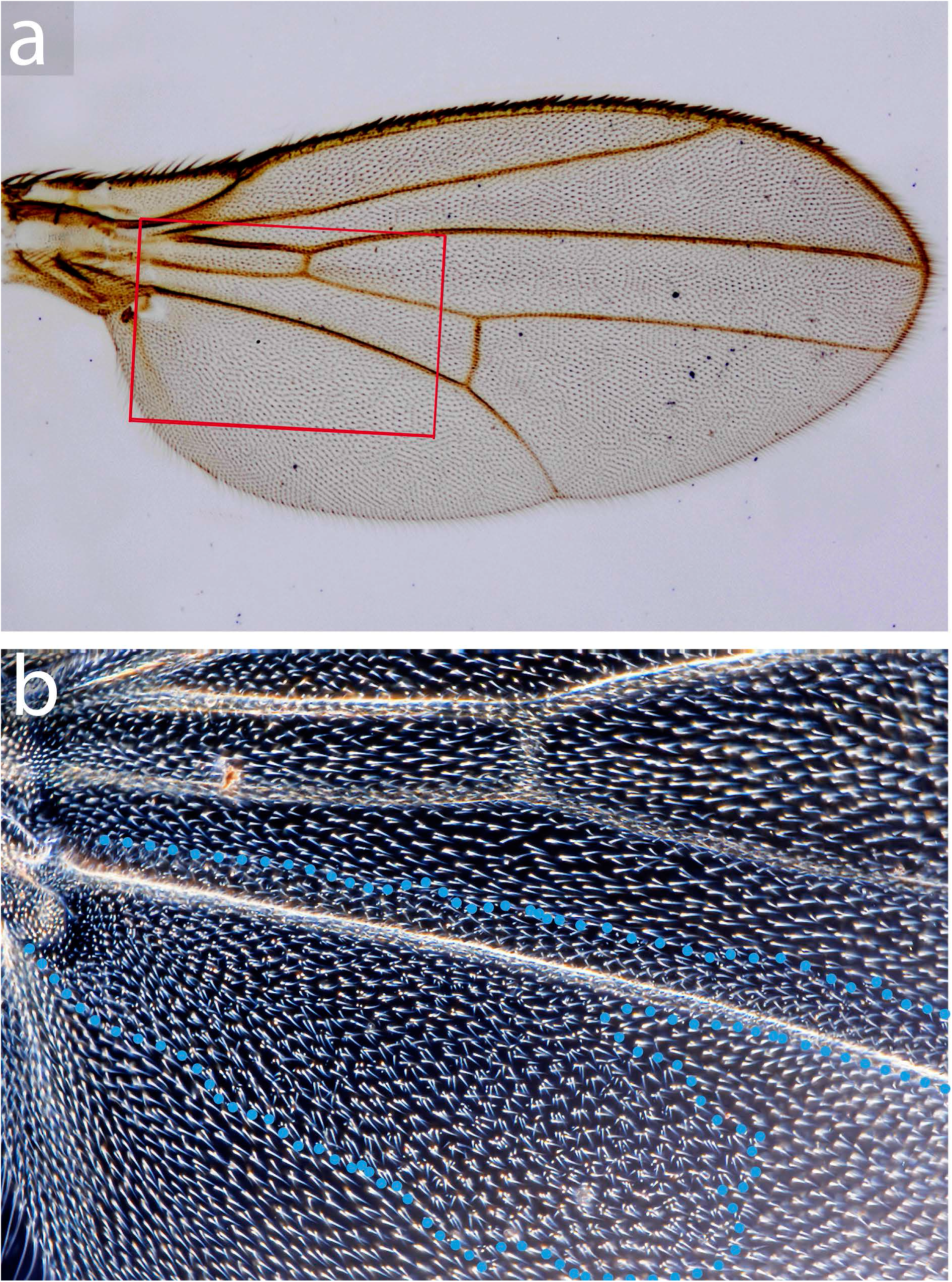
A clone marked with *mwh* that crosses the putative F-P border near vein L5. This is a single clone (outlined in blue dots) confined to the dorsal wing surface, it was made in a *mwh*/+ fly by X-irradiation at 36 ± 12 hrs. **A** the wing (bright field microscopy) **B** the clone (dark field). In these experiments the clone frequency is low (Table 1) so the pattern shown is very unlikely to be due to two or more independent clones (p < 0.0001). There are many other clones like this one that cross well over vein L5 (Table 1).

The data shown in Bryant ^29^ and in Gonzalez-Gaitan et al ^33^ also clearly refute an F-P compartment border, their figures show several clones crossing the putative F-P boundary at different stages. Moreover we have examined a large number of unpublished drawings of more clones made by one of us (JFdC). These latter clones were marked in a slightly different way (twin clones marked either with *mwh* or *f*^36*a*^) but again they deny the existence of the F-P border, clones of both genotypes frequently crossing from one side to the other (Table 1). Again we see a tendency for clones to run parallel to part of vein L5 for stretches, this behaviour occurs mainly proximal to the posterior crossvein. However distal to this crossvein most clones cross the L5 vein and, generally, the tendency for clonal boundaries to follow veins is intermittent and not as precisely defined as a compartment border. It can be emphasised again that both *f*^36*a*^ and *mwh* are mutations that are homozygous viable with no significant effect on wing size, shape or pattern.

Figure 2 in Abbasi and Marcus ^22^ suggests the F-P border might lie a few cells posterior to vein L5, but even this line is not respected by our clones, neither the ones made with a growth advantage, nor without. We find this to be true both for clones mostly occupying territory between veins L4 and L5, and for clones located mostly between L5 and the posterior wing margin.

**Figure 2.**
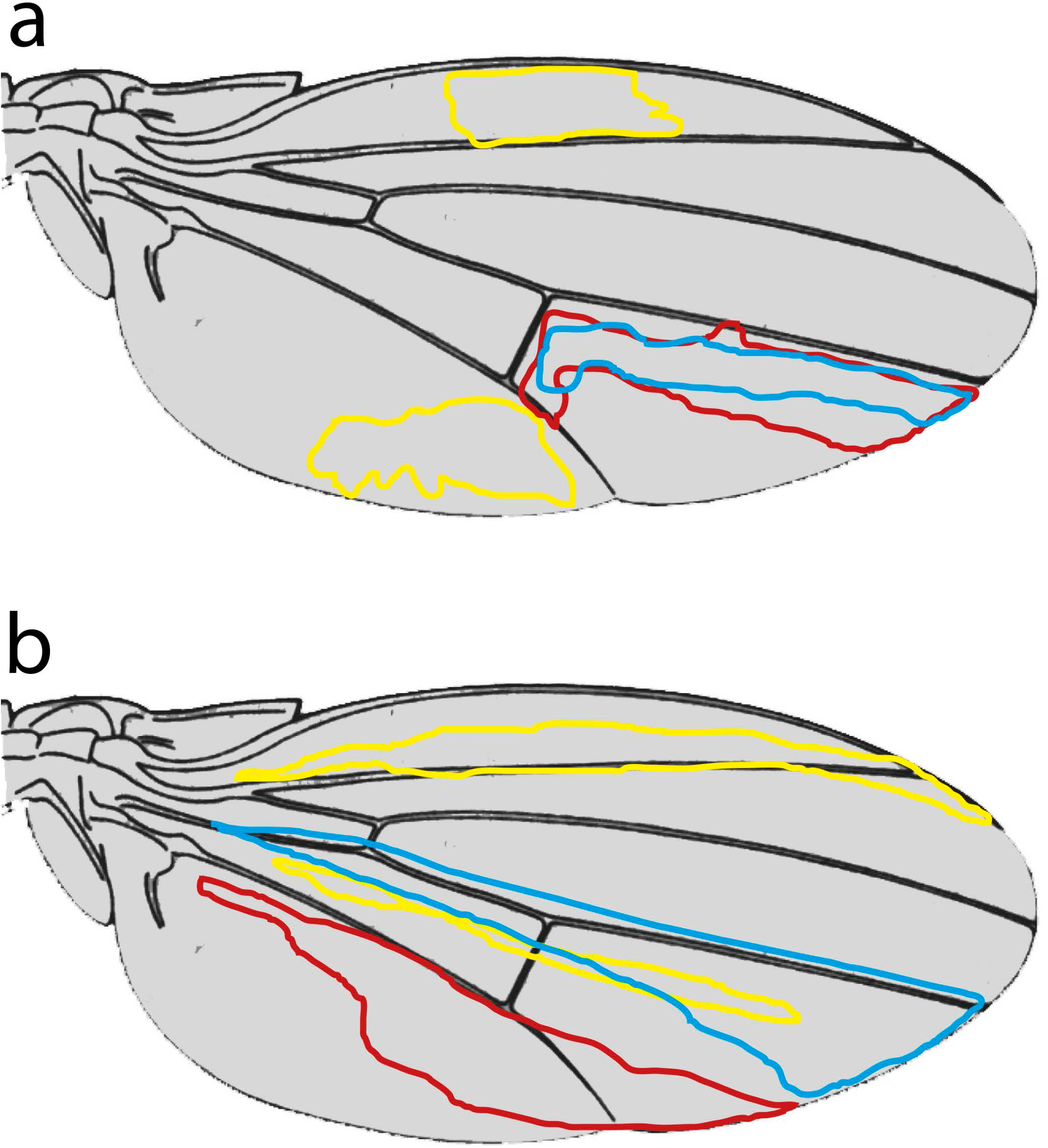
Wings with outlines of clones. **A** shows clones taken from Figure 2 in Abbasi and Marcus ^22^ that we find implausible because we have never found clones shaped like these. **B** shows typical clones that occupy the same areas drawn by JFdC from a large library of such clones.

We have six additional concerns about the clones as reported in Abbasi and Marcus ^22^:

**First**, in our hands, *yellow* (*y*^1^), the marker used, is not reliable. We could not clearly define the clonal boundaries with *y*^1^ and we did not employ it as a marker of the wing blade. We have found *yellow* to be an excellent marker for bristles ^34^ but to be inadequate for the wing surface; we can detect large patches in the wing blade only when both dorsal and ventral wing surfaces carry the mutation.
**Second**, it is strange that Abbasi and Marcus ^22^ do not mention that their clones are mostly confined to one wing surface (dorsal or ventral), as they must be (see Figure 1). Indeed we think detecting and precise mapping of *y*^1^ clones that are restricted to one surface would be demanding. Our worries could be allayed by a photograph showing that a *y*^1^ clone confined to the dorsal or ventral wing surface can be delineated precisely. In this context it is surprising that their Figure 2a shows yellow clones all with non overlapping territories as we would suppose that some are dorsal and some ventral and therefore would frequently be found overlaying each other.
**Third**, some of the clone shapes reported by Abbasi and Marcus ^22^ are implausible to us (JFdC) as we have never found such shapes in our preparations (ca 300 clones), nor in any of the shapes shown in the figures of Bryant ^29^ and González-Gaitán et al. ^33^. Particularly relevant are the clones in the region between L4 and L5 ending or starting near the posterior crossvein and all clones posterior to L5. In the first case clones do not respect the crossvein and tend to be more elongated than shown in their Figure 2, while clones distal to the crossvein to the L5 vein tend to cross this vein in a majority of cases (see our Figure 2). A similar discrepancy is also observed with their clone outlines located near the L2 vein, which in our hands tend to be much more elongated than square (see our Figure 2 for examples).
**Fourth**, again concerning Figure 2 in Abbasi and Marcus ^22^, this Figure shows some clones crossing the anteroposterior border very substantially and these clones are implausible. Abbasi and Marcus ^22^ point this out and suggest it might be due to a defect in the method used to superimpose clonal shapes on a standardised wing. They argue that “geometric distortion in the anterior of the wing images partially obscures the well-studied A-P boundary that is also present”. However this explanation does not cover the two pink-outlined clones in their Figure 2, b1 which cross the A-P boundary for long distances. An explanation here might be that each shape-outline is produced by two independent clones on one wing; but at first sight this is not likely because of the low frequency of their clones (but see **Fifth**, below).
**Fifth** We are concerned by the low frequency of clones reported by Abbasi and Marcus ^22^: 44 clones in 1778 wings. In standard flippase experiments it is usual to obtain several clones per wing. Possibly Abbasi and Marcus are missing many *y*^1^ clones, which if true would again question the validity of the *yellow* marker in the wing blade. Furthermore, their Figure 2a suggests that some wings carry several clones which, if so, presumably means that most of the 1778 wings had no clones, and a small minority had several, which is odd.
**Sixth** Their Figure 2 shows only 6 clones that approach the F-P border from the anterior side, as against many more from the posterior side (we cannot count them exactly but there appears to be 20 or more from the posterior side). We would expect that, in an unbiased sample, approximately equal numbers of clones would approach a legitimate compartment boundary from each side. However, we believe *yellow* clones would be easier to spot away from the wing centre, if so that could explain this discrepancy but raises a further question about the validity of the *yellow* marker.

### A posterior organizer in the Drosophila wing?

Although our evidence argues strongly against the existence of an F-P lineage boundary some evidence does point to a secondary organiser within the posterior compartment of the *Drosophila* wing. There is a site of *dpp* expression in a small proximal region of the posterior compartment. This source generates a local gradient of Dpp expression and is required for the normal pattern of the wing; in its absence the alula and much of the axillary cord are missing ^35^. Still, we cannot completely exclude the possibility that, during evolution, dipterans such as *Drosophila* might have lost a F-P lineage boundary but kept a late expression of Dpp expression in the posterior wing.

## Materials and Methods

Larvae of genotypes (JFdC) *Dp(3;Y;1)M2, f*^36*a*^; *mwh* or *sha*/*ck*, (GM/PAL) *M*(*3*)*i^55^*/ *mwh jv* or +/ *mwh jv* were irradiated with a 1000R dose at the indicated AEL (after egg laying) times. Wings with *mwh*/*f* or *sha*/*ck* twin clones or *mwh jv* clones were mounted in GMM medium and analysed under the microscope.

## Contributions

PAL wrote the main manuscript text and JC prepared the figures. PAL, JFdC and GM designed and obtained experimental data. All authors reviewed the manuscript.

## Acknowledgements

Work in our laboratories is funded by grants BFU2015-64220-P (JFdC) and BFU-2015-67839-P (GM) from the Spanish Ministerio de Economia y Competitividad, and 107060/Z/15/Z (PAL) from the Wellcome Trust.

This paper was submitted to *Scientific Reports* in January 2018 and favourably reviewed in the spring. Since then publication has been awaiting the response of the original authors of the study we are criticising. In the continuing absence of this response we decided to place this preprint in bioRxiv so that people can read it.

## Data Availability

The datasets generated during and/or analysed during the current study are available from the corresponding author on reasonable request.

